# Defects in lipid homeostasis reflect the function of TANGO2 in Acyl-CoA metabolism

**DOI:** 10.1101/2022.11.05.515282

**Authors:** Agustin Lujan, Ombretta Foresti, Nathalie Brouwers, Alex Mateo Farre, Alessio Vignoli, Jose Wojnacki, Vivek Malhotra

## Abstract

We show that TANGO2, which lacks a transmembrane domain localizes predominantly to mitochondria and transiently to endoplasmic reticulum (ER) and lipid droplets (LDs). Evaluation of lipids in HepG2 cells lacking TANGO2 revealed an increase in the size of lipid droplets and reactive oxygen species production. There is also a marked increase lysophosphatidic acid (LPA) and a concomitant decrease in its biosynthetic precursor phosphatidic acid (PA). These changes are exacerbated in nutrient starved cells. Based on our data, we suggest that the principle function of TANGO2 is in acyl-CoA metabolism, which is necessary for the acylation of LPA to generate PA. This defect subsequently affects metabolism of many other fatty acids. These data help explain the physiological consequence of TANGO2 that induce acute metabolic crisis including rhabdomyolysis, cardiomyopathy and cardiac arrhythmias often leading to fatality upon starvation and stress.

## Introduction

In 2006, we reported a collection of new genes that were required for transport and organization of the Golgi complex in drosophila ^1^.

TANGO2, a gene unrelated to TANGO1, which lacks a transmembrane domain is shown to be cytosolic and also localized to the mitochondria. TANGO2 is reported to functions both in endoplasmic reticulum (ER) to Golgi transport and in mitochondrial physiology, however, its precise role is not known ^2^. Mutations in TANGO2 result in metabolic encephalopathy and arrhythmias and these are exacerbated in conditions of nutrient starvation leading to fatality ^3,4^. What is the physiological role of TANGO2 and how mutations in TANGO2 lead to fatality in conditions of starvation?

We show here that TANGO2 in mammalian cells is localized predominantly to mitochondria, but also partially at sites of ER and lipid droplets (LDs) juxtaposed to mitochondria. Knockdown of TANGO2 increased the size of LDs and elevated intracellular reactive oxygen species (ROS) levels. Mass spectrometry of cellular lipid composition revealed that TANGO2 lacking cells exhibit markedly high lyso-phosphatidic acid (LPA) and low levels of phosphatidic acid (PA). These cells are also highly reduced in cardiolipin (CL), which is ordinarily produced from PA. Interestingly, changes in these cellular properties are exacerbated in TANGO2 lacking cells cultured in nutrient free media. Based on our data, we propose that TANGO2 is functions mainly in lipid homeostasis at the level acyl-CoA metabolism.

## Results

### TANGO2 localizes predominantly to the mitochondria, but also transiently to LDs and ER

TANGO2 in humans has six isoforms produced as a result of alternative splicing. The TANGO2-1(TNG2_HUMAN·Q6ICL3-1) and TANGO2-2 (TNG2_HUMAN·Q6ICL3-2) isoforms are the most similar to their orthologs, such as hgr-9 in worms, YGR127W in yeast, and tango2 in zebrafish (Figure 1 A-B). It is important to note that the antibody used to TANGO2 recognizes both isoforms in human cells and the tagged versions of both isoforms show the same location (Figure 1 A). The siRNA based depletion reduces levels of all isoforms of TANGO2. In the following sections, the data on the location pertains to isoform 1 and the effect of TANGO2 depletion on cell physiology is a result of reduction of all isoforms.

**Figure 1.**
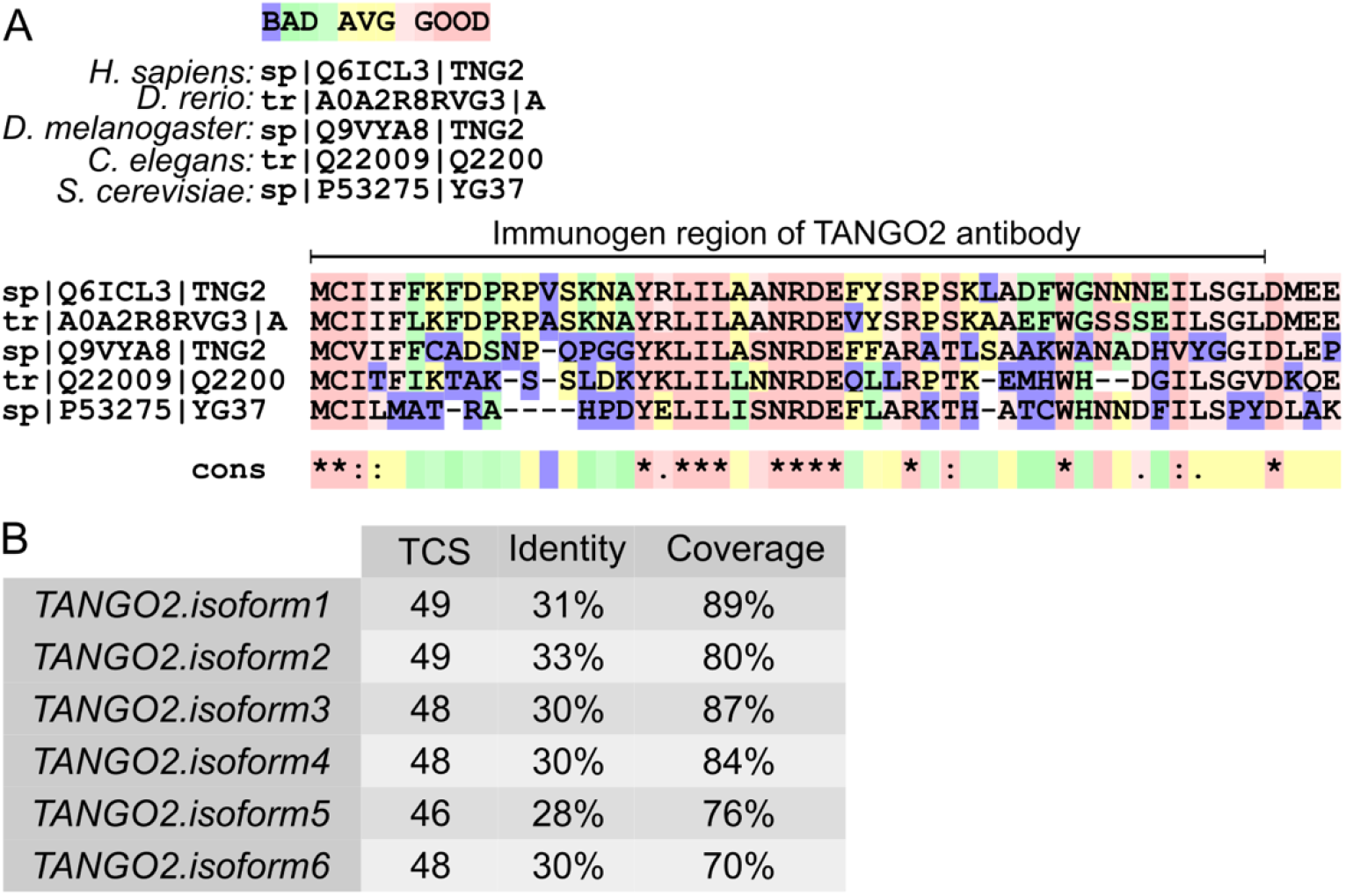
TANGO2 orthologs. **A)** In silico sequence alignment of TANGO2 orthologs in human (*H. sapiens*), zebrafish (*D. rerio*), fruit fly (*D. melanogaster*), worm (*C. elegans*), and yeast (*S. cerevisiae*) using T-COFFEE software. **B)** Multiple sequence alignment (MSA) of each human TANGO2 isoform with species orthologs. The transitive consistency score (TCS) identify the most correct alignment positions in a MSA, the Identity evaluate the sequence function, and the Coverage describes the average number of reads that align to known reference bases.

Previous published data revealed the location of TANGO2 to mitochondria in cells and in mitochondrial enriched membranes ^2,5^. But this has also been challenged ^6^. We monitored the location of TANGO2 distribution dynamically in living cells. We generated mScarlet and EGFP-tagged at the C-terminus of TANGO2. We co-overexpressed, by transient transfections, this form of TANGO2 with ER (ER-pmTurquoise2)-, Mitochondrial (Mitochondria-pmTurquoise2)-, Peroxisomal (Peroxisome-SKL-mTurquoise2)-, or LDs (GPAT4-mScarlet)-fluorescent specific plasmids in HepG2 cells. All images were acquired by live-cell confocal and time-lapse microscopy to avoid fixation artifacts. We observed high co-localization (Pearson coefficient, r=0.88) between TANGO2 and the mitochondrial marker. TANGO2 was also found at sites of mitochondria closely juxtaposed to ER and LDs (Figure 2). To avoid potential artifacts from an uneven or abnormal protein expression by transient transfection, we generated EGFP-tagged at the C-terminus of the TANGO2 using lentiviruses by the standard procedures ^7^. TANGO2 distribution in HepG2 cells, by this procedure, was analogous to that observed in cells expressing tagged TANGO2 by transient-transfection (Figure S1 A). In addition, there was also a diffuse cytosolic staining of TANGO2 (Figure S1 B). To check whether cleavage of EGFP from TANGO2 might have occurred in cells expressing tagged TANGO2, we verified the size of expressed construct by SDS/PAGE followed by western blotting with an anti-GFP antibody (Figure S1 C). The size of the construct expressed matches the expected molecular weight of an EGFP tagged TANGO2.

**Figure 2.**
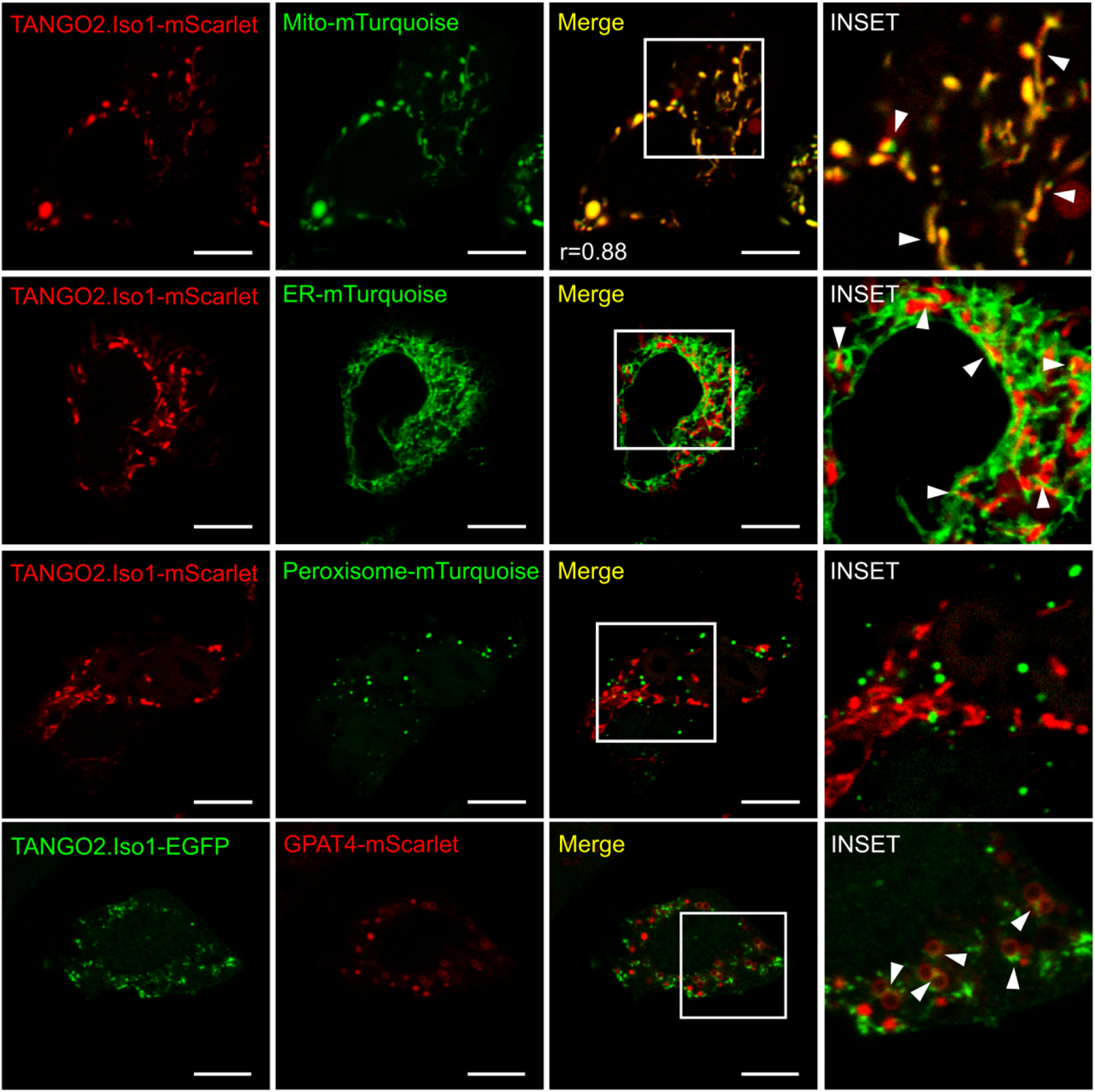
TANGO2 location in HepG2 cells. Cells expressing TANGO2.Iso1-mScarlet or TANGO2.Iso1-EGFP co-transfected with Mitochondria-pmTurquoise2, ER-pmTurquoise2, Peroxisome-SKL-mTurquoise2 or GPAT4-mScarlet to detect lipid droplets. Arrowheads indicate colocalization between two labels. Pearson coefficients (r) was calculated with coloc2 plugin in ImageJ software. Images are representative of three independent experiments. Scale bars = 10 µm.

### TANGO2 lacking cells contain larger Lipid droplets (LDs)

As shown above TANGO2 frequently appears at sites that are marked by LDs. This prompted us to test if cells lacking TANGO2 affect any aspect of LDs. We transiently silenced TANGO2 (siTANGO2) by using specific siRNA. A control was generated with non-targeting siRNA as described henceforth as control or Mock. The silencing procedure reduced TANGO2 levels by ∼40% after one round of siRNA transfection. However, after 2 sequential rounds of transfection a ∼70% reduction in TANGO2 levels was evident (Figure S2 A). We have used the latter scheme of double transfection to reduce TANGO2 levels in all experiments presented below (Figure S2 B). Mock HepG2 cells and TANGO2 depleted cells were cultured in medium containing 10% Fetal Bovine Serum (FBS) (DMEM complete) or without FBS (starvation) for 4 h. The respective cells were then incubated with membrane permeant neutral lipid marker HCS LipidTOX™ Deep Red (red) and the DNA marker Hoechst-33342 (blue) for 30 min at 37 ºC followed by live-cell imaging. LDs were consistently large in TANGO2 depleted cells compared to mock cells (Figure 3 A, left panel). Moreover, nutrient deprived TANGO2 depleted cells (Starvation; Figure 3 A, right panel) had even larger LDs. To confirm that these structures are indeed LDs, TANGO2 depleted HepG2 cells were fixed and visualized with an antibody to adipose differentiation-related protein that specifically localizes to LDs (ADRP; red). Nuclear DNA in these cells was labeled with DAPI (blue). There was indeed an increase in the size of ADRP containing LDs upon depletion of TANGO2 (Figure 3B). To quantify this increased retention of neutral lipids upon depletion of TANGO2 visualized by confocal microscopy, we developed a macro for Image J Software. This procedure analyzes Z-stack images to determine the number and volume of each LipidTOX-positive particle randomly. Five samples were used for each condition, with a total of 67 control cells and 75 TANGO2 depleted cells. As shown in Figure 3, total volume of LDs per cell (panel C) and the volume of each LD (panel D) was significantly higher in TANGO2 depleted compared to the Mock cells (*p*=0.0005 and *p*=0.002, respectively). However, the number of LDs in each cell remain the same regardless of the TANGO2 levels (Figure 3 E). The increase in size, therefore, is not by fusion of smaller LDs under these experimental conditions. To further strengthen our proposal on the effect of TANGO2 on LD size, we added 120 nm Oleic acid (OA) at 37 º C for 6 h to both mock and TANGO2 depleted cells. Changes in the total volume of LDs was then evaluated by fluorescence intensity (FI) of HCS LipidTOX with flow cytometry (Figure 3F). Three independent experiments were performed for each condition in triplicates. Loss of TANGO2 revealed a significant increase of total LDs volume compared to Mock-cells (Figure 3 G).

**Figure 3.**
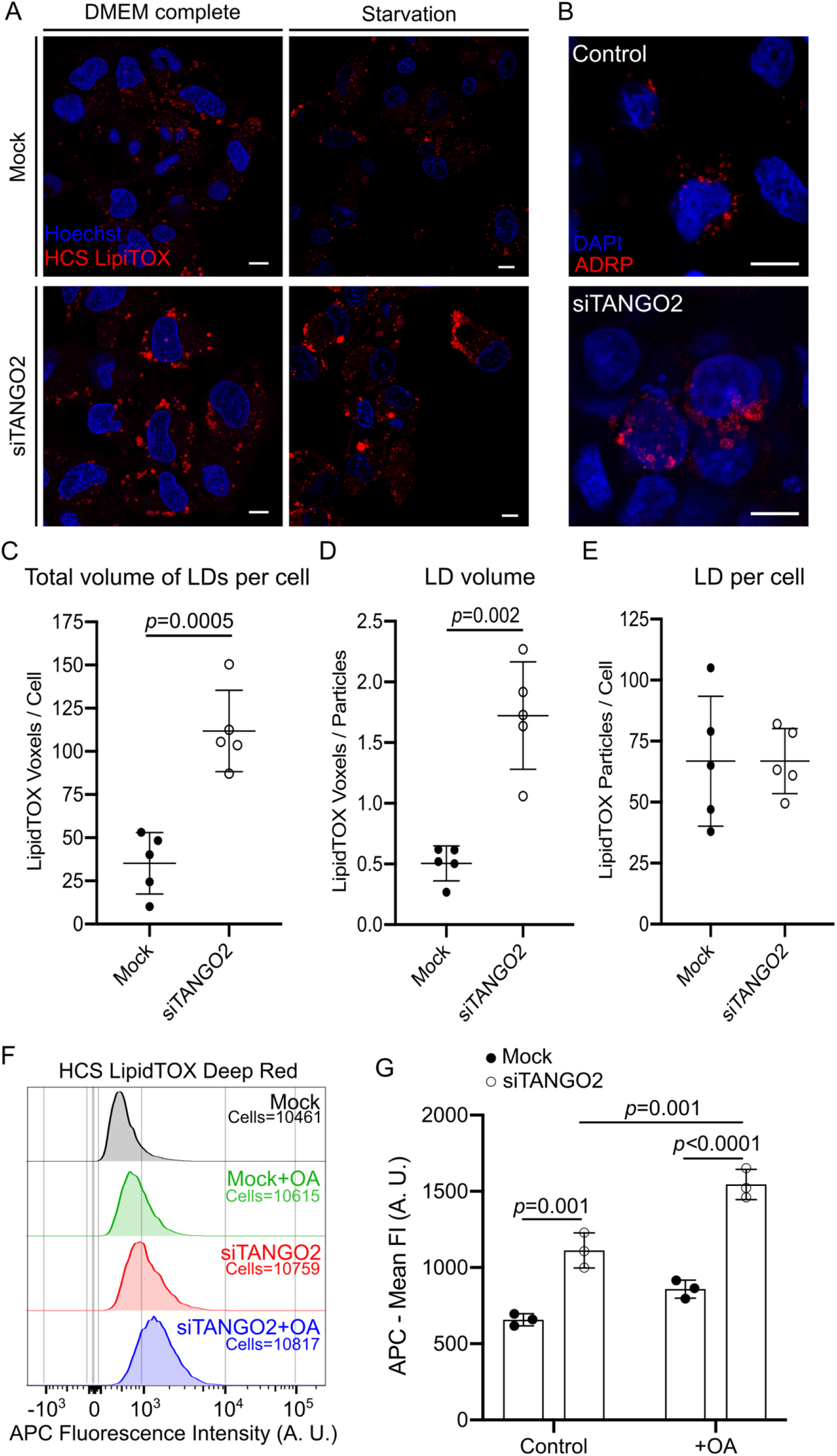
TANGO2 depletion affect the size of LDs. **A)** Confocal images of HCS LipidTOX marker (red) and Hoechst-33342 (blue) in mock (top) and siTANGO2 (bottom) HepG2 cells cultured in control conditions (DMEM complete; left panel) or starved of nutrients (starvation; right panel). Scale bars = 10 µm. **B)** Mock (top) and siTANGO2 (bottom) cells in control conditions were fixed and incubated with anti-ADRP (red) and DAPI (blue). Scale bars = 10 µm. **C-E)** Quantification of LDs in mock (n=67 cells) and TANGO2 depleted (n=75 cells) cells in five samples for each condition by an ImageJ macro developed in our lab (open code in GitHub. **C)** Total volume of LDs present per cell. **D)** Volume of each LD in a cell. **E)** Number of LDs per cell. **F)** Mock and siTANGO2 HepG2 cells maintained in control conditions or with 120 nM oleic acid (OA), and incubated with HCS LipidTOX Deep Red for 30 min before flow cytometry analysis. Neutral lipids were detected by measuring the fluorescence intensity of APC. **G)** Mean fluorescence intensity of APC in three different samples of each condition. Images and graphs are representative of three independent experiments. In graphs, boxes and bars are the mean ± SD.

### TANGO2 loss affects the quantities of specific lipids

The change in LD size in TANGO2 depleted cells prompted us to analyze changes in the cellular lipid composition. It is also important to note that patients with TANGO2 deficiency suffer metabolic crises upon fasting, which suggests its potential function in lipid metabolism. Three independent preparation of mock cells, TANGO2 depleted cells, starved mock cells and starved TANGO2 depleted cells were analyzed for their total lipid composition (Lipotype GmbH). There was no significant difference in the total amount of lipid between TANGO2 silenced groups compared to control for each condition. The mole percent fraction (molp) quantification allows comparison between samples, disregarding the absolute amount of lipids.

We then analyzed changes in quantities of lipids by comparing each lipid species in mock samples under control and starvation conditions (Figure 4, left panel); mock and TANGO2 depleted (Figure 4, middle panel), and starved TANGO2 depleted compared with mock starved cells (Figure 4, right panel). In starved mock cells lysophosphatidic acid (LPA) levels are low, whereas lysophosphoglycerol (LPG), phosphoglycerol (PG) and phosphatidic acid (PA) levels are notably high (Figure 3 left panel). In control cells LPA is used to produce PA, which is also a precursor of PG and mitochondria specific cardiolipin (CL). Comparison of lipids in TANGO2 depleted cells to mock revealed an increase in levels of LPA and reduction in PA and CL levels. There was also an increase in most lyso-phospholipids and a reduction in the corresponding phospholipids (middle panel). Importantly, in starving TANGO2 depleted cells, there was a further increase in LPA and a decrease in PA levels (Figure 4, right panel). It is important to note that the levels of LPG and PG are high in mock cells cultured in starvation medium (left panel). In TANGO2 depleted cells, LPG levels are reduced and there is a dramatic reduction in PG levels. The levels of these two lipids are further reduced in starving TANGO2 cells. We suggest these lipids are used to produce CL in starving TANGO2 depleted cells as evident by the slight recovery in its levels under these conditions.

**Figure 4.**
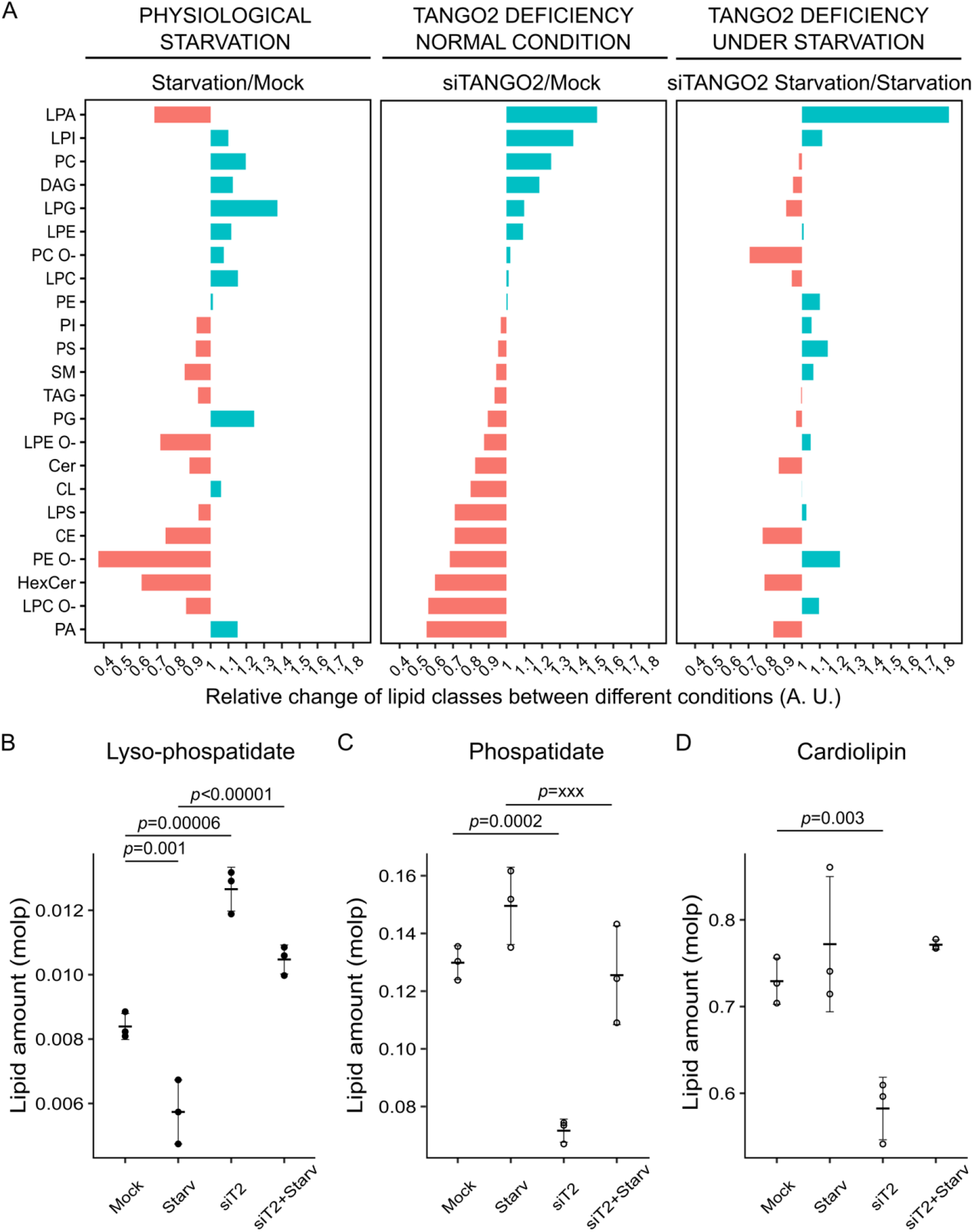
Lipidomics of TANGO2 depleted cells for comparison with control and nutrient starved cells. **A)** Relative change of lipid class according to different conditions: physiological starvation (left panel), TANGO2 depleted cells in control conditions (middle panel), and TANGO2 depleted cells upon nutrient starvation (right panel). **B)** Quantification of LPA changes in mock and siTANGO2 depleted cells expressed in mole percent fractions. **C)** Quantification of PA changes in mock and siTANGO2 depleted cells expressed in mole percent fractions. **D)** Quantification of CL changes in mock and siTANGO2 depleted cells expressed in mole percent fractions. Data is representative of three independent experiments. In graphs, boxes are the mean ± SD.

These data show that nutrient-starved TANGO2-depleted cells are severely affected in LPA to PA ratio and there is a defect in additional fatty acids homeostasis.

### TANGO2 depleted cells exhibit high levels of reactive oxygen species (ROS)

Mitochondria produce ROS at the inner mitochondrial membrane during oxidative phosphorylation. ROS produced as a byproduct of ATP synthesis is useful for cells, but high levels of ROS unless scavenged are toxic. The fact that TANGO2 localizes to mitochondria and an earlier report of an increased in mitochondrial-ROS in cells from TANGO2 deficiency patients ^5^ prompted us to test this feature in our experimental conditions. TANGO2 depleted cells with 10% FBS (DMEM complete), or without FBS (Starvation), and mock cells were incubated for 20 min with CellROX Deep Red reagent to measure cellular ROS levels and MitoTracker Green to measure total mitochondrial mass by flow cytometry analysis. Cell viability was measured with using CellTrace Calcein Green AM and DAPI to demonstrate that TANGO2 silencing did not induce cell death in our samples (Figure S3). To determine the total mitochondrial mass, we analyzed the FI of FITC for all samples (Figure 5 A). The statistical analysis shows the mean FI (MFI) of FITC in three independent experiments. The results reveal no significant difference in total mitochondrial mass in TANGO2 depleted cells compared to controls (Figure 5 B).

**Figure 5.**
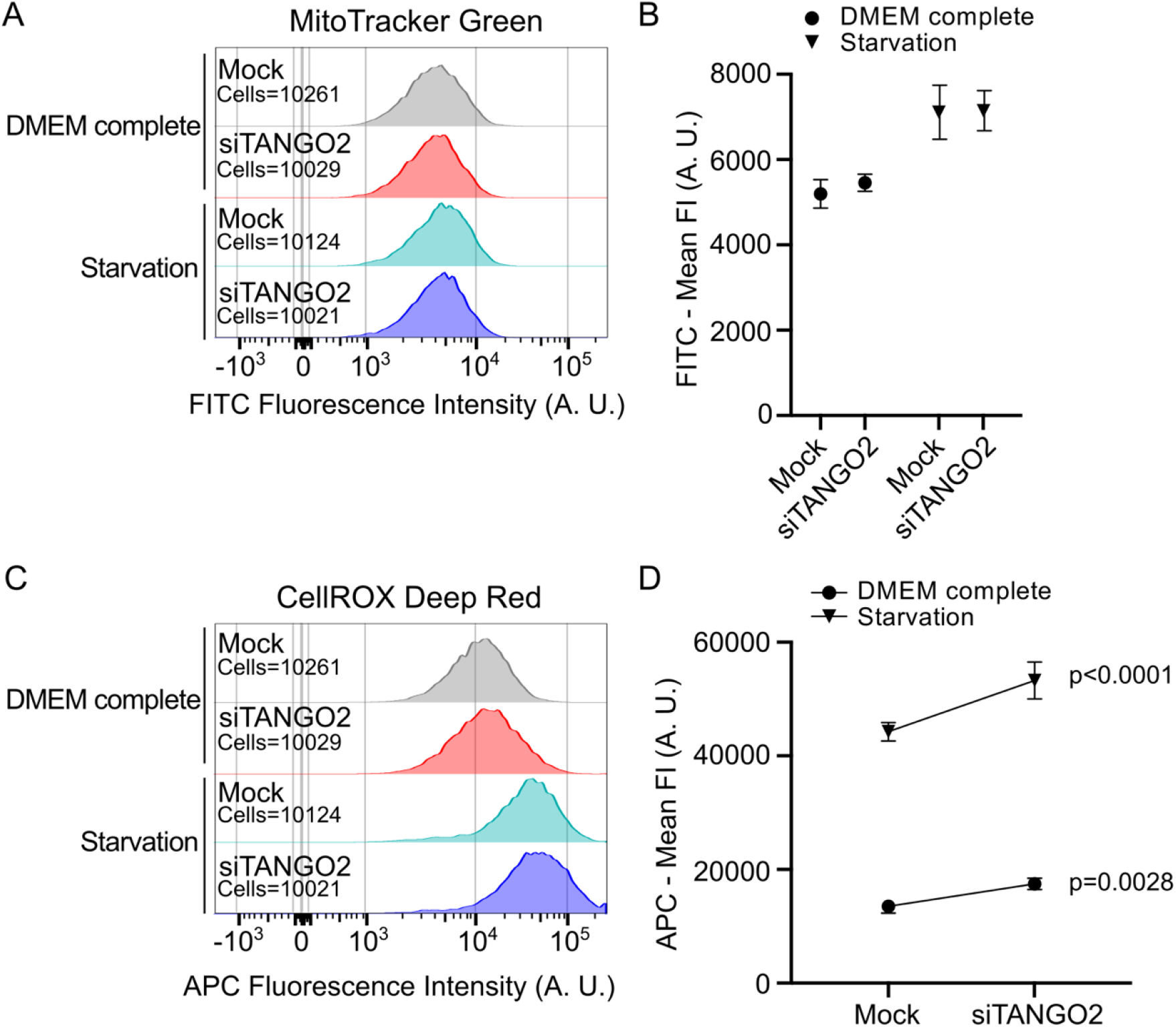
TANGO2 depleted cells exhibit increased ROS levels. **A-D)** Mock and siTANGO2 cells were incubated in control conditions (DMEM complete) or in nutrient lacking medium (starvation) for 4 h. Cells were incubated with CellROX Deep Red to detect ROS levels and MitoTracker Green to monitor mitochondrial mass. **A)** Mitochondrial mass was detected by measuring the fluorescence intensity of FITC by flow cytometry. **B)** The mean fluorescence intensity of FITC was measured in three different samples of each condition. **C)** Cellular ROS level was detected by measuring the fluorescence intensity of APC by flow cytometry. **D)** The mean fluorescence intensity of APC was measure in three different samples of each condition.

To measure ROS production in these experimental conditions, we analyzed the difference in FI of APC corresponding to CellROX marker (Figure 5 C). The data show that depletion of TANGO2 significantly increases ROS levels, which is further elevated in conditions of nutrient starvation (Figure 5 D).

## Discussion

Identification of TANGO genes for their involvement in Transport and Golgi Organization is revealing unexpected insights into cell physiology ^1^. Studies on TANGO1 has revealed how cells organize ER exit sites to collect and export cargoes that are too big for secretion by a standard COPII vesicle of 60 nm average diameter ^8,9^. Importantly, the function of TANGO1 fits well with the physiology and associated pathologies ^10,11^. Other genes characterized from this collection are shown to be required for cargo sorting at the level of the Golgi apparatus ^12^ and for trafficking of a new class of transport containers called CARTS that move secretory cargo from Golgi apparatus to the cell surface ^13^. TANGO2, a less studied protein from the TANGO collection, is gaining attention recently because of its connection to human pathologies ^14–17^. Patients with mutations in TANGO2 exhibit defects in cardiac and neuronal physiology leading to fatalities upon starvation and stress (https://tango2research.org) ^3,18^. Here we describe its function and its link to starvation induced pathologies in patients with mutations in TANGO2.

### TANGO2 functions in lipid homeostasis

TANGO2 has been shown to function at the mitochondria, and in the ER to Golgi step of protein secretion pathway ^2^. The role of TANGO2 in ER to Golgi pathway of secretion fits well with its original assigned function in transport and Golgi organization ^1^. Our new data reveal that TANGO2 localizes predominantly to mitochondria, but also at sites on mitochondria that are juxtaposed to LDs, and ER.

Loss of TANGO2 in HepG2 cells has revealed four clear features. 1, a change in the ratio of LPA to PA. LPA levels are elevated and PA is reduced. 2, reduction in CL content. 3, increase in the size of LDs. 4, increase in ROS level. The significance of these changes to TANGO2 function in physiology and pathology follows.

1. LPA is acylated to generate PA. The fact that LPA increases and PA decreases suggests that acylation of LPA is defective in TANGO2 lacking cells. The enzymes (LPAATs) involved in these acylation reactions are transmembrane protein and acyl-CoA produced in the cytosol has to be imported into the lumen by membrane embedded transporters. TANGO2 lacks a transmembrane domain and cannot therefore function as an enzyme or a membrane inserted acyl-coA importer to catalyze these reactions in the lumen or the membrane of the ER. We suggest it likely functions on the cytoplasmic face of mitochondria and the ER in the synthesis of acyl-CoA or for its collection for import into the lumen of these compartments. Our data show that although predominantly located to mitochondria, there is an enrichment of TANGO2 at sites juxtaposed to ER. This could be the site of LPA to PA conversion. We suggest that without TANGO2, acyl-CoA is unavailable for the synthesis of PA from LPA. As a result, LPA levels rise and there is a concomitant drop in PA. This effect is exacerbated in nutrient starved TANGO2 depleted cells
2. PA is used for the production of CL at the inner mitochondrial membrane. A decrease in PA levels depleted cells explains the reduced amounts of CL in TANGO2 cells. This would also affect mitochondrial physiology. In starving TANGO2 depleted cells, CL levels recover slightly compared to TANGO2 depleted cells. We suggest that when PA is low, cells likely use PG to produce CL. This might explain reduction in PG levels in starving TANGO2 depleted cells compared to TANGO2 cells cultured in nutrient containing medium. Changes in CL are associated to a number of cardiac diseases ^19–22^. For example, Barth syndrome (BTHS) is an inherited cardiomyopathy, associated with skeletal myopathy, growth retardation and neutropenia ^23–25^. BTHS is caused by a mutation in a mitochondrial phospholipid-lysophospholipidtransacylase, involved in the biogenesis of CL ^26,27^. These features are similar to the pathologies associated with TANGO2 mutations. \
3. The change in LDs, we suggest is due to changes in the levels of acyl-coA and LD specific fatty acids that utilize this metabolite for their growth. Analysis of the lipid droplet composition will help address this outcome in cells depleted of TANGO2. The increase in ROS levels suggests that mitochondria are not defective in ATP production *per se*.

Clearly, TANGO2 depleted cells exhibit a change in lipid metabolism, which is aggravated in cells starving cells

### TANGO2 as a haem transfer protein

A paper published recently by Chen and colleagues shows that loss of human TANGO2 and its orthologs in yeast and worms causes accumulation of haem in compartments like mitochondria and lysosome related organelles (LROs) ^28^. The authors show that adding recombinant TANGO2 to mitochondria enriched membrane fractions in vitro stimulates haem transfer to apo-myoglobin and recombinant TANGO2 has low affinity binding to ferrous and ferric haem. It is concluded that haem is exported from the lumen into the cytoplasm by transmembrane transporters where it is captured by TANGO2 and subsequently transferred to specific clients. This is observed even in trypsinised mitochondria and the authors conclude that cytoplasmic facing domains of transmembrane proteins or attached proteins are not required in TANGO2 mediated haem transfer. Why TANGO2 depleted cells accumulate haem in mitochondria (or LROs) in the first place? If TANGO2 captures haem post its export from mitochondria, as is reported even in trypsinised mitochondria, then haem should be distributed into the cytoplasm and not retained in mitochondria or LROs. The data also do not explain the aggravated effects of TANGO2 depletion linked to human pathologies including lack of any obvious sign of anaemia as would be evident for defects in haem physiology. A more plausible explanation for the data is that amphipathic haem is retained in mitochondria because of altered membrane lipid composition upon TANGO2 depletion. Further work is therefore required to clarify whether the proposed function of TANGO2 in haem transfer from mitochondria might be a consequence of dysfunctional mitochondrial lipid composition.

## Conclusion

We propose that TANGO2 has a function in events leading to acylation of LPA to produce PA. Sacher and colleagues have reconstituted many features of TANGO2 mutations in flies ^29^. Interestingly, most of the aberrations in fly physiology are corrected, as well as an ER to Golgi transport defect in human cells, by vitamin B5 supplement, which is used for the synthesis of CoA. In other words, loss of TANGO2 function is restored by replenishing or overdosing events leading to CoA synthesis. CoA is used for acyl-CoA production, which is then used to acylate LPA to form PA. This fits well with our data: LPA increases and PA decreases in TANGO2 depleted cells and this change is exacerbated upon starvation. A defect in PA levels impacts the synthesis of mitochondrial CL. In order to balance this deficiency, the cells use other potential substrates such as the PG to produce CL. The net effect is an imbalance in lipid composition or homeostasis that we suggest also increases ROS levels. The acyl-coA pathway is also used for the synthesis of numerous fatty acids and a defect in its production would change the form and function LDs to influence cellular lipid homeostasis. The starving cells rely heavily on lipids for the ATP production and TANGO2 depleted are thus further stressed. These findings explain the role of TANGO2 and how mutations trigger a variety of cardiac, muscular and neurological pathologies leading to fatalities upon conditions of starvation and stress.

## Acknowledgments

We thank all members of the Malhotra laboratory for valuable discussions and critical reading of the manuscript. We thank Aida Rodriguez for advice with ROS analysis. We acknowledge the support of the Spanish Ministry of Science, the Centro de Excelencia Severo Ochoa, and the CERCA Programme / Generalitat de Catalunya. V. Malhotra is an Institució Catalana de Recerca i Estudis Avançats professor at the Centre for Genomic Regulation. V. Malhotra is an Institució Catalana de Recerca i Estudis Avançats professor at the Centre for Genomic Regulation Work in the Malhotra lab is funded by grants from the Spanish Ministry of Economy and Competitiveness (Plan Nacional to VM: PID2019-105518GB-I00) and the European Research Council Synergy Grant (ERC-2020-SyG - Proposal No. 951146). AL is funded by the European Molecular Biology Organization (EMBO ALTF 659-2021), JW is funded by the European Research Council (H2020-MSCA-IF-2019-894115). OF is a Ramon y Cajal fellow in the Malhotra lab (RYC-2016-20919). This work reflects only the authors’ views, and the EU Community is not liable for any use that may be made of the information contained therein.

## Materials and Methods

### Cell culture

HepG2 cells were cultured in Dulbecco’s Modified Eagle’s Medium (DMEM) cell culture media (Lonza) supplemented with 10% (vol / vol) heat-inactivated fetal bovine serum (FBS; Gibco) (DMEM complete), 100 units/mL penicillin, and 100 μg/mL streptomycin (Gibco), at 37 °C in a humidified incubator supplied with 5% CO2. For starvation conditions, cells were incubated in DMEM without FBS for 4 h.

### Plasmids, RNA interference, and cell transfection

For transient protein over-expression, cells were plated and 24 h later transfected with 1:1 ratio (DNA plasmid:Lipofectamine) using Lipofectamine 3000 reagent (ThermoFisher Scientific) following manufacturer recommendations. Plasmids generated in our lab: TANGO2.Isoform1-EGFP, TANGO2.Isoform1-mScarlet, Peroxisome-SKL-mTurquoise2, GPAT4-mScarlet. Plasmids used and available from Addgene: ER-pmTurquoise2 (Addgene #36204), Mitochondria-pmTurquoise2 (Addgene #36208). For TANGO2 silencing, cells were plated and transfected with 1:3 ratio (RNA:Lipofectamine) using Lipofectamine RNAiMAX reagent (ThermoFisher Scientific) following manufacturer recommendations. 24 h after the first transfection, cells were transfected with a second and identical mix (ratio 1:3) of RNA:Lipofectamine. Two days after the second transfection, experiments described in this paper with siTANGO2 conditions were performed. TANGO2 RNAi sequences: Hs.Ri.TANGO2.13.1-SEQ1 and Hs.Ri.TANGO2.13.1-SEQ2 (IDT technologies).

### Antibodies and Reagents

Primary antibodies used: anti-ADRP (Santa Cruz), anti-GFP (Abcam), anti-β-actin (Abcam). Secondary antibodies used: Alexa Fluor 594 donkey antirabbit (Invitrogen), Alexa Fluor 647 donkey anti-rabbit (Invitrogen), Reagents used were: DAPI (Invitrogen), Hoechst-33342 (Sigma), HCS LipidTOX Deep Red (ThermoFisher Scientific), Mitotracker Green (ThermoFisher Scientific), CellROX Deep Red (ThermoFisher Scientific), Calcein AM (ThermoFisher Scientific), FluorSave reagent (CalBiochem), Lipofectamine RNAiMAX reagent (ThermoFisher Scientific), Lipofectamine 3000 reagent (ThermoFisher Scientific).

### Immunoblotting

Immunoblot analysis was performed following the Biorad general protocol recommendations. Briefly, equal amounts of protein were resolved by SDS/ PAGE, transferred onto 0.45-µm PVDF membranes (Amersham), and incubated overnight with anti-GFP (1:1000; Abcam), anti-β-actin (1:1,0000; Sigma) antibodies followed by Alexa-conjugated secondary antibodies (1:10,000; Invitrogen) for 1 h at 37 ºC. Bands were visualized an Odyssey Clx (LI-COR Biosciences).

### Immunofluorescence and Immunohistochemistry

HepG2 cells (1 × 10^5^ cells per well) were grown on coverslips. After mock or siTANGO2 silencing, cells were fixed with 3% paraformaldehyde, quenched, and incubated overnight with anti-ADRP (1:1000) at 4 ºC followed by Alexa Fluor 647 donkey anti-rabbit (1:2000) and DAPI (1:10,000) for 1 h at 37 ºC. Coverslips were washed and mounted in FluorSave reagent. Samples were analyzed and high resolution images were acquired in an inverted Leica TCS SP8 confocal microscope equipped with photomultipliers and hybrid detectors. Images were processed using ImageJ software.

### Live cell imaging

HepG2 cells (2.5 × 10^5^ cells per dish) were grown on 35 mm polymer-bottom dishes (ibidi). After mock or siTANGO2 silencing, cells were washed and the cell medium was replaced with the appropriate conditioned medium for the indicated period of time. The cells were then washed several times to remove the conditioned medium and incubated with the appropriate reagent (Hoechst-33342, HCS LipiTOX Deep Red, MitoTracker Green). Finally, after several washes, cells were used to perform the experiments described in the paper. Samples were analyzed at 37 ºC in 5% CO2 atmosphere. High resolution and time-lapse images were acquired in an inverted Leica STELLARIS confocal microscope equipped with photomultipliers and hybrid detectors. Images were processed using ImageJ software. For co-localization analysis, we calculated the Pearson Correlation Coefficient of confocal images with the coloc2 plugin in ImageJ software. To quantify the volume of LDs in confocal images, we developed a macro (open source code in www.github.com/AAMateo/Lipid_Droplets) to identify each voxel (pixel^3^) stained by the lipophilic marker LipidTOX in Z-stacks. We then selected five Z-stacks from each condition. This macro automatically identified and distinguished the positive from the negative particles on each slide and generated a data set with the volume of each particle, the number of particles, and the total volume of positive particles. This method reduced the chances in counting errors or biases.

### Flow cytometry assays

To examine the total LDs volume, HepG2 cells (1×10^5^) were seeded in 24-well plates (ThermoFisher Scientific). Two days after the second silencing (Mock or siTANGO2), cells were washed and the cell medium was changed with the appropriate medium for the times indicated. The cells were then washed several times to remove the conditioned medium and incubated with the appropriate reagent (CellROX Deep Red, HCS LipidTOX Deep Red, MitoTracker Green, Calcein AM, DAPI). Cells were gently detached using trypsin and washed several times before flow cytometry analyses. Flow cytometry assays were performed by technical triplicates of at least 10,000 cells per condition each time in three independent experiments. In the statistical model these triplicates are nested together. Flow cytometry analysis was performed using a BD LSR II Flow Cytometer (Becton Dickinson Biosciences).

### Lipid extraction for mass spectrometry lipidomics

Mass spectrometry-based lipid analysis was performed by Lipotype GmbH (Dresden, Germany) as described (Surma et al. 2021). Lipids were extracted using a chloroform/methanol procedure (Ejsing et al. 2009). Samples were spiked with internal lipid standard mixture containing: cardiolipin 14:0/14:0/14:0/14:0 (CL), ceramide 18:1;2/17:0 (Cer), diacylglycerol 17:0/17:0 (DAG), hexosylceramide 18:1;2/12:0 (HexCer), lyso-phosphatidate 17:0 (LPA), lyso-phosphatidylcholine 12:0 (LPC), lyso-phosphatidylethanolamine 17:1 (LPE), lyso-phosphatidylglycerol 17:1 (LPG), lyso-phosphatidylinositol 17:1 (LPI), lyso-phosphatidylserine 17:1 (LPS), phosphatidate 17:0/17:0 (PA), phosphatidylcholine 17:0/17:0 (PC), phosphatidylethanolamine 17:0/17:0 (PE), phosphatidylglycerol 17:0/17:0 (PG), phosphatidylinositol 16:0/16:0 (PI), phosphatidylserine 17:0/17:0 (PS), cholesterol ester 20:0 (CE), sphingomyelin 18:1;2/12:0;0 (SM), triacylglycerol 17:0/17:0/17:0 (TAG). After extraction, the organic phase was transferred to an infusion plate and dried in a speed vacuum concentrator. The dry extract was re-suspended in 7.5 mM ammonium formiate in chloroform/methanol/propanol (1:2:4, V:V:V). All liquid handling steps were performed using Hamilton Robotics STARlet robotic platform with the Anti Droplet Control feature for organic solvents pipetting.

### MS data acquisition

Samples were analyzed by direct infusion on a QExactive mass spectrometer (Thermo Scientific) equipped with a TriVersa NanoMate ion source (Advion Biosciences). Samples were analyzed in both positive and negative ion modes with a resolution of Rm/z=200=280000 for MS and Rm/z=200=17500 for MSMS experiments, in a single acquisition. MSMS was triggered by an inclusion list encompassing corresponding MS mass ranges scanned in 1 Da increments (Surma et al. 2015). Both MS and MSMS data were combined to monitor CE, DAG and TAG ions as ammonium adducts; LPC, LPC O-, PC, PC O-, as formiate adducts; and CL, LPS, PA, PE, PE O-, PG, PI and PS as deprotonated anions. MS only was used to monitor LPA, LPE, LPE O-, LPG and LPI as deprotonated anions; Cer, HexCer and SM as formiate adducts.

### Lipidomics data analysis and post-processing

Data were analyzed with an in-house developed lipid identification software based on LipidXplorer (Herzog et al. 2011; Herzog et al. 2012). Data post-processing and normalization were performed using an in-house developed data management system. Only lipid identifications with a signal-to-noise ratio >5, and a signal intensity 5-fold higher than in corresponding blank samples were considered for further data analysis.

### Statistical analysis

Statistical analysis was performed using GraphPad Prism (GraphPad) and R softwares. Data represent the mean ± SDs of N experiments. For multiple comparisons, ANOVA with subsequent Bonferroni’s or Tukey’s posttests was used. P values less than 0.05 were considered statistically significant.

## Figures

**Figure S 1.**
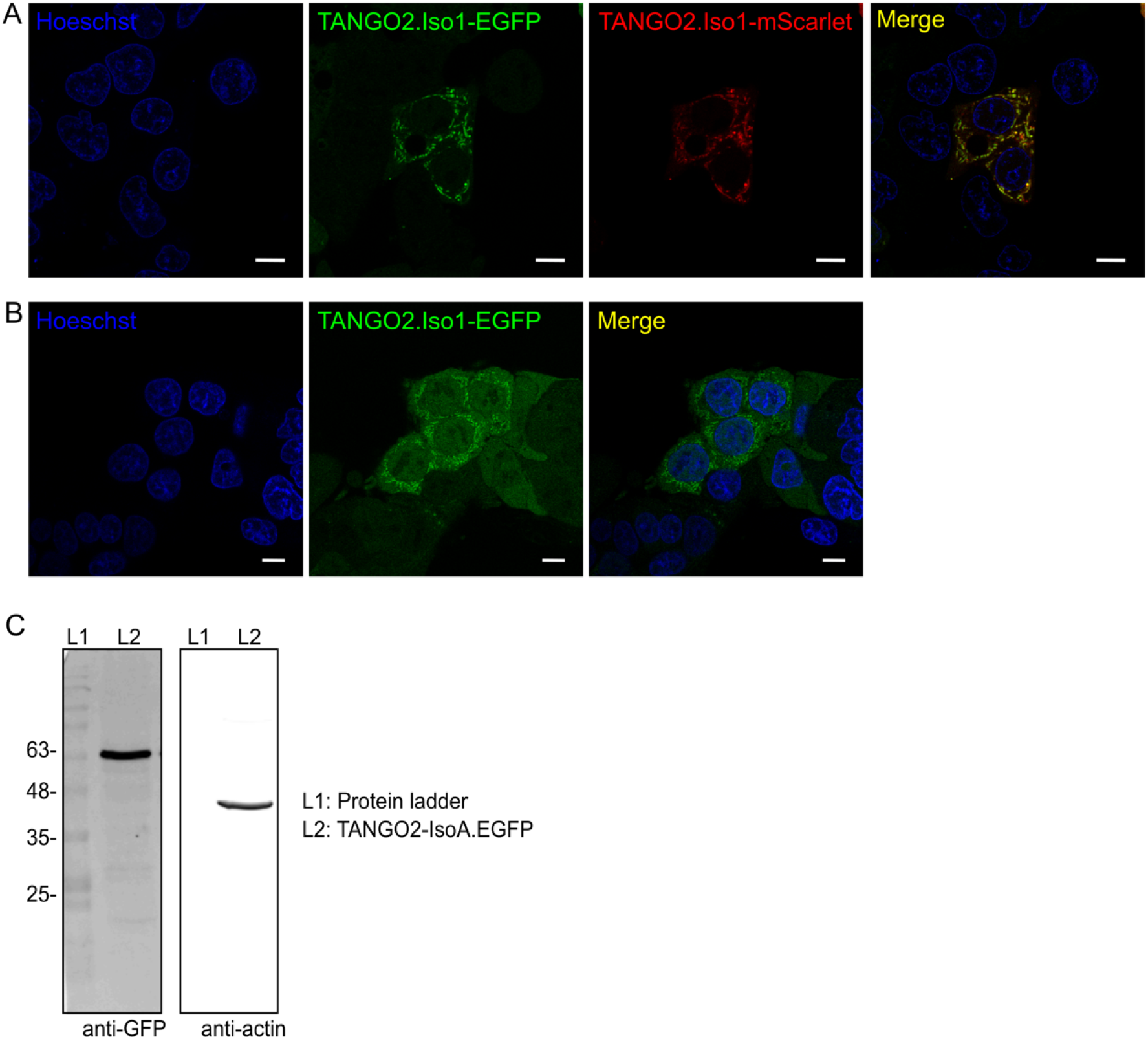
TANGO2-tagged analogous. **A)** TANGO2.Iso1-EGFP HepG2 stable cells were transiently transfected with TANGO2.Iso1-mScarlet (red), and labeled with Hoechst (blue). Merge shows all overlapping channels. **B)** TANGO2.Iso1-EGFP (green) HepG2 stable cell was labeled with Hoechst (blue). **C)** Immunoblot analysis of HepG2_TANGO2.Iso1-EGFP stable cell line incubated with anti-GFP and anti-actin. Anti-actin was used as loading control.

**Figure S 2.**
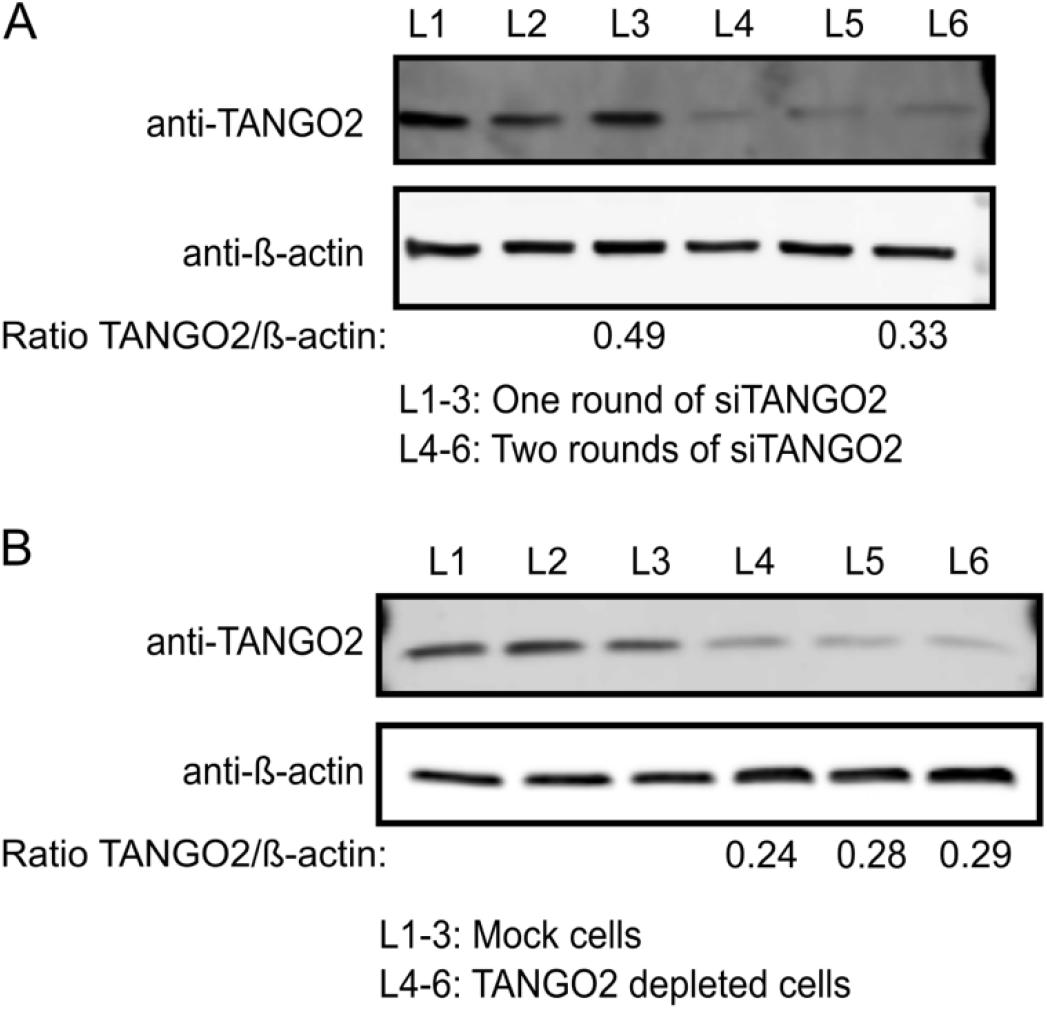
TANGO2 depleted cells. **A-B)** Immunoblot analysis of HepG2 wild type cells silenced by specific TANGO2 siRNA. **A)** Immunoblot analysis comparison between one or two rounds of TANGO2 silencing. **B)** Immunoblot analysis of two rounds of TANGO2 silencing compared with non-targeting siRNA (mock).

**Figure S 3.**
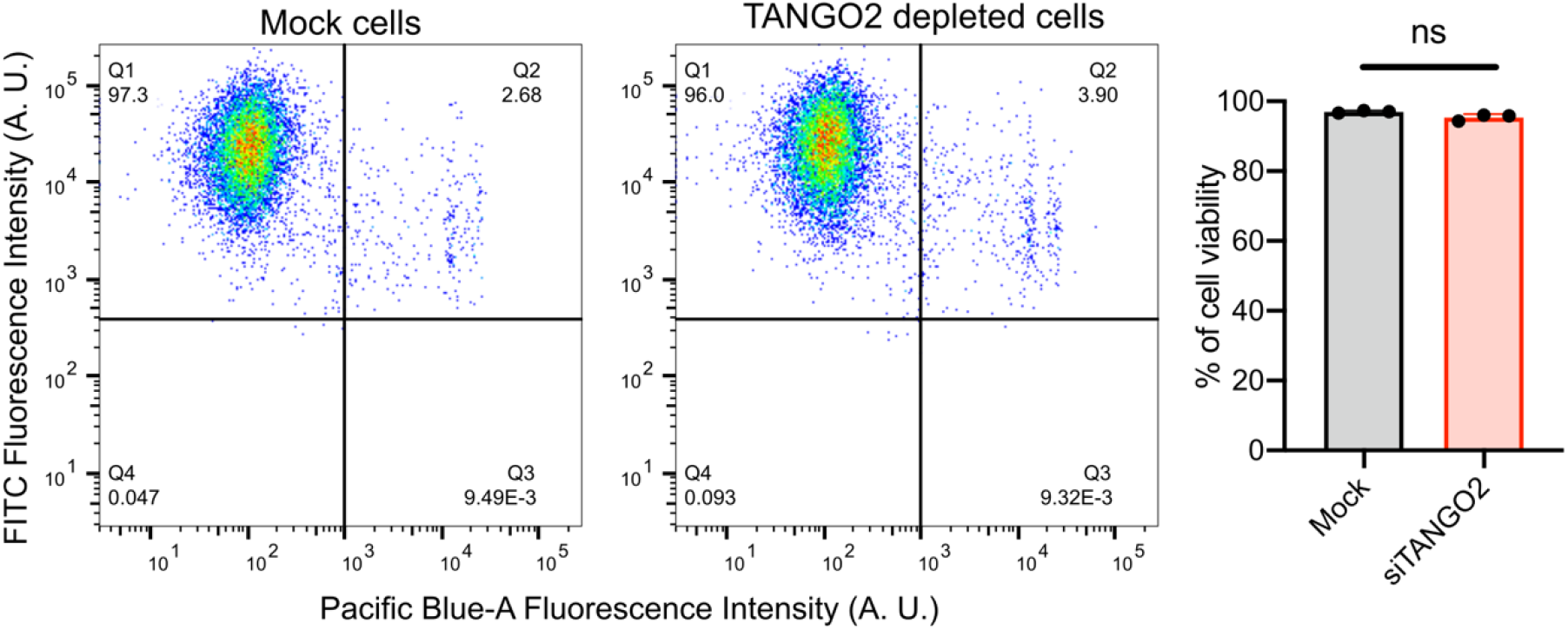
Cell viability in TANGO2 depleted cells. Mock and TANGO2 depleted cells were incubated with CellTrace Calcein Green AM (FITC) and DAPI (Pacific Blue-A) to determine cell viability by flow cytometry. Each dot in the column graph represent one of three independent experiments.

## References

1. Bard, F. et al. Functional genomics reveals genes involved in protein secretion and Golgi organization. Nature 439, 604–607 (2006).

2. Milev, M. P. et al. The phenotype associated with variants in TANGO2 may be explained by a dual role of the protein in ER-to-Golgi transport and at the mitochondria. J. Inherit. Metab. Dis. 44, 426–437 (2021).

3. Lalani, S. R. et al. Recurrent Muscle Weakness with Rhabdomyolysis, Metabolic Crises, and Cardiac Arrhythmia Due to Bi-allelic TANGO2 Mutations. Am. J. Hum. Genet. 98, 347–357 (2016).

4. Kremer, L. S. et al. Bi-allelic Truncating Mutations in TANGO2 Cause Infancy-Onset Recurrent Metabolic Crises with Encephalocardiomyopathy. Am. J. Hum. Genet. 98, 358–362 (2016).

5. Heiman, P. et al. Mitochondrial dysfunction associated with TANGO2 deficiency. Sci. Rep. 12, 3045 (2022).

6. Jennions, E. et al. TANGO2 deficiency as a cause of neurodevelopmental delay with indirect effects on mitochondrial energy metabolism. J. Inherit. Metab. Dis. 42, 898–908 (2019).

7. Tandon, N., Thakkar, K. N., LaGory, E. L., Liu, Y. & Giaccia, A. J. Generation of Stable Expression Mammalian Cell Lines Using Lentivirus. Bio-protocol 8, (2018).

8. Saito, K. et al. TANGO1 facilitates cargo loading at endoplasmic reticulum exit sites. Cell 136, 891–902 (2009).

9. Raote, I. & Malhotra, V. Tunnels for Protein Export from the Endoplasmic Reticulum. Annu. Rev. Biochem. 90, 605–630 (2021).

10. Lekszas, C. et al. Biallelic TANGO1 mutations cause a novel syndromal disease due to hampered cellular collagen secretion. Elife 9, (2020).

11. Guillemyn, B., Nampoothiri, S., Syx, D., Malfait, F. & Symoens, S. Loss of TANGO1 Leads to Absence of Bone Mineralization. JBMR plus 5, e10451 (2021).

12. von Blume, J. et al. Actin remodeling by ADF/cofilin is required for cargo sorting at the trans-Golgi network. J. Cell Biol. 187, 1055–1069 (2009).

13. Wakana, Y. et al. A new class of carriers that transport selective cargo from the trans Golgi network to the cell surface. EMBO J. 31, 3976–3990 (2012).

14. Schymick, J. et al. Variable clinical severity in TANGO2 deficiency: Case series and literature review. Am. J. Med. Genet. A 188, 473–487 (2022).

15. Powell, A. R., Ames, E. G., Knierbein, E. N., Hannibal, M. C. & Mackenzie, S. J. Symptom Prevalence and Genotype-Phenotype Correlations in Patients With TANGO2-Related Metabolic Encephalopathy and Arrhythmias (TRMEA). Pediatr. Neurol. 119, 34–39 (2021).

16. Hoebeke, C., Cano, A., De Lonlay, P. & Chabrol, B. Clinical phenotype associated with TANGO2 gene mutation. Arch. Pediatr. 28, 80–86 (2021).

17. Bérat, C.-M. et al. Clinical and biological characterization of 20 patients with TANGO2 deficiency indicates novel triggers of metabolic crises and no primary energetic defect. J. Inherit. Metab. Dis. 44, 415–425 (2021).

18. Miyake, C. Y. et al. Cardiac crises: Cardiac arrhythmias and cardiomyopathy during TANGO2 deficiency related metabolic crises. Hear. Rhythm 19, 1673–1681 (2022).

19. Mulligan, C. M., Le, C. H., deMooy, A. B., Nelson, C. B. & Chicco, A. J. Inhibition of delta-6 desaturase reverses cardiolipin remodeling and prevents contractile dysfunction in the aged mouse heart without altering mitochondrial respiratory function. J. Gerontol. A. Biol. Sci. Med. Sci. 69, 799–809 (2014).

20. Paradies, G. et al. Decrease in mitochondrial complex I activity in ischemic/reperfused rat heart: involvement of reactive oxygen species and cardiolipin. Circ. Res. 94, 53–59 (2004).

21. Petrosillo, G. et al. Mitochondrial dysfunction associated with cardiac ischemia/reperfusion can be attenuated by oxygen tension control. Role of oxygen-free radicals and cardiolipin. Biochim. Biophys. Acta 1710, 78–86 (2005).

22. Dudek, J. Role of Cardiolipin in Mitochondrial Signaling Pathways. Front. cell Dev. Biol. 5, 90 (2017).

23. Barth, P. G. et al. X-linked cardioskeletal myopathy and neutropenia (Barth syndrome): respiratory-chain abnormalities in cultured fibroblasts. J. Inherit. Metab. Dis. 19, 157–160 (1996).

24. Cantlay, A. M. et al. Genetic analysis of the G4.5 gene in families with suspected Barth syndrome. J. Pediatr. 135, 311–315 (1999).

25. Steward, C. G. et al. Neutropenia in Barth syndrome: characteristics, risks, and management. Curr. Opin. Hematol. 26, 6–15 (2019).

26. Bione, S. et al. A novel X-linked gene, G4.5. is responsible for Barth syndrome. Nat. Genet. 12, 385–389 (1996).

27. Dudek, J. & Maack, C. Barth syndrome cardiomyopathy. Cardiovasc. Res. 113, 399–410 (2017).

28. Sun, F. et al. HRG-9 homologues regulate haem trafficking from haem-enriched compartments. Nature 610, 768–774 (2022).

29. Asadi, P., Milev, M. P., Saint-Dic, D., Gamberi, C. & Sacher, M. Vitamin B5, a Coenzyme A precursor, rescues TANGO2 deficiency disease associated defects in Drosophila and human cells. bioRxiv 2022.11.04.514597 (2022). doi:10.1101/2022.11.04.514597

